# Early exposure to high-fat diet impairs central and peripheral metabolic function: Impacts on cognition and mitochondrial function

**DOI:** 10.1101/2022.06.22.496258

**Authors:** Wembley Rodrigues Vilela, Paula Maria Quaglio Bellozi, Victor Luna Picolo, Bruna Neves Cavadas, Keila Valentina Silva Marques, Louise Tavares Garcia Pereira, Angélica Amorim Amato, Kelly Grace Magalhães, Márcia Renata Mortari, Jair Trapé Goulart, Andreza Fabro de Bem

## Abstract

The impact of overnutrition early in life is not restricted to the onset of cardiovascular and metabolic diseases, but also affects critical brain functions related to cognition. This study aimed to evaluate the relationship between peripheral metabolic and bioenergetic changes induced by high-fat diet (HFD) and their impact on hippocampal cognitive functions in juvenile mice. To this purpose, three-week-old male C57BL/6 mice received a HFD or control diet for seven weeks, associated with two low doses of streptozotocin (STZ) or vehicle, to accelerate the metabolic dysfunction. HFD induced metabolic changes in mice, particularly related to glucose metabolism, in spite of the absence of obesity and changes in lipid profile. HFD exposure starting from weaning impaired recognition and spatial memories in mice, without inducing a depressive-like behavior. Increased immunoreactivity for GFAP and a trend towards a decrease in NeuN staining were verified in the hippocampus of HFD-fed mice. HFD caused a bioenergetic impairment in the hippocampus, characterized by a decrease in both O_2_ consumption related to ATP production and in the maximum respiratory capacity. The thermogenic capacity of brown adipose tissue was impaired by HFD, here verified through the absence of a decrease in O_2_ consumption after UCP-1 inhibition and increase in the reserve respiratory capacity. Impaired mitochondria function was also observed in the liver of HFD mice, while no changes were verified in O_2_ consumption in the heart of juvenile mice. These results indicate that the introduction of a HFD early in life has a detrimental impact on bioenergetic and mitochondrial function of tissues with metabolic and thermogenic activities, which is likely related to hippocampal metabolic changes and cognitive impairment.

**Highlights:**

- HFD introduced early in life impacts mitochondrial function
- Dietary shift early in life leads hippocampal dysfunction
- Early life HFD exposure disrupts BAT thermogenic acitivity
- HFD-induced hippocampal and BAT mitochondrial dysfunction impacts cognition

## 1. INTRODUCTION

Twenty first century children are being precociously exposed to a Western lifestyle, characterized by physical inactivity and ingestion of ready-to-eat ultraprocessed foods, rich in refined carbohydrates and fats (Di Cesare et al., 2019; Malone and Hansen, 2019; WHO, 2020). This dietetic shift puts children in the path of metabolic derangements, such as obesity, type 2 diabetes mellitus and hypertension, that were once considered adult problems. Alongside the fact that an overweight child strongly tends to become an obese adolescent and adult, critical brain functions are particularly affected after early exposure to a Westernized diet (Cunningham et al., 2014; Yang et al., 2019). This is worrisome, since childhood is a period of neurobehavioral shaping, required for long-life cognitive processing (Spear, 2000).

Clinical evidences have shown the impact of overweight and obesity on children’s cognitive function. A 31-year follow-up study evidenced a direct correlation between cardiovascular risk factors and obesity in childhood with cognitive impairment in adulthood. The longitudinal increase in systolic blood pressure, total cholesterol and body weight was associated with worse cognitive scores in midlife, described as alterations in episodic memory, associative learning and sustained attention (Hakala et al., 2021). Bauer and colleagues reported that overweight and obesity in children reduced hippocampal volume and impaired executive cognitive performance (Bauer et al., 2015). Overweight children had worse scores in spatial cognitive tasks on the mental rotation test (Jansen et al., 2011). Moreover, increased body fat in children was associated with poorer cognitive performance in planning, attention, math and reading (Davis and Cooper, 2011).

Energy-dense diet protocols in animals (high fat, high carbohydrate and high sucrose) have been largely used to mimic human Western lifestyle (excessive consumption of processed foods and sweetened beverages) as well as to understand biologic mechanisms involved in metabolic derangements (de Bem et al., 2020; Picolo et al., 2021; Trevino et al., 2015). Not only metabolic active peripheral tissues such as liver (Ishimoto et al., 2013), white adipose tissue (WAT) (Politis-Barber et al., 2020) and brown adipose tissue (BAT) (Gao et al., 2020) are affected by the consumption of these diets, but also, and particularly the brain is susceptible to structural and functional damages (Trevino et al., 2017; Yang et al., 2019).

Recent preclinical studies in rodents have shown that a high-fat diet (HFD) introduced early in life impaired hippocampal functions. Two different studies showed that hippocampal-dependent contextual fear and spatial memory impairment were associated with diminished long-term potentiation in CA1 hippocampal slices in juvenile mice exposed to HFD (Khazen et al., 2019; Yang et al., 2019). A decreased neurogenic ability in the dentate gyrus (DG), as well as increased microgliosis and proinflammatory cytokines levels were observed in hippocampus from mice exposed to HFD early in life (Vinuesa et al., 2019; Vinuesa et al., 2016). Interestingly, HFD impaired neurogenic ability and relational memory in juvenile, but not adult mice (Boitard et al., 2012), suggesting that adolescence represents a vulnerable period when cognitive functions can be strongly modulated by the diet.

Mitochondria is a key organelle globally affected by HFD consumption, where lipid overload causes structural and functional changes (Aoun et al., 2012; Putti et al., 2015). It is know that mitochondrial dysfunction contributes to the pathogenesis of metabolic disorders (Pinti et al., 2019). Recent studies from our and other groups suggest that the hippocampal and BAT mitochondria are particularly sensible to HFD intake. Yang and colleagues observed that the excess of lipid intake from diet diminishes mitochondrial biogenesis and alters fusion-fission protein levels in hippocampus of mice (Yang et al., 2021). We also verified a reduced hippocampal mitochondrial function in adult mice fed a HFD (de Paula et al., 2021). Furthermore, BAT mitochondrial metabolism has currently gathered attention as a therapeutic target for obesity and related metabolic disorders (Lettieri Barbato et al., 2015; Tseng et al., 2010). Recent evidence suggests that the thermogenic activity of BAT was impaired in offspring pups exposed to HFD during pregnancy and lactation (Lettieri Barbato et al., 2015). Considering that affected tissues include not only those participating in nutrient metabolism, such as adipose, liver and skeletal muscle, but also the brain, we hypothesize that a global and connected bioenergetic crisis could drive the behavioral changes induced by HFD in juvenile mice. Our results show that HFD consumption early in life leads to metabolic dysfunction and cognitive impairment in early adulthood, suggesting that mitochondria are placed as the hub that crosstalk the brain and periphery.

## 2. MATERIAL AND METHODS

### Animals and Experimental design

Juvenile C57BL/6 male mice (postnatal day 21, P21; immediately after weaning) were housed in acrylic cages with filtered air system (Alesco; 3-4 mice/cage), in controlled temperature (23-25°C), and 12h light/dark cycle (lights on 6 a.m.) with access to food and water *ad libitum*. Mice were fed with a standard rodent diet (CD; 15% fat) or a high-fat diet (HFD; 48% fat) (both from PRAGSOLUÇÕES Biociências®) for seven weeks (see complete nutritional composition on Table S1). To boost the metabolic changes, two consecutive intraperitoneal injections of streptozotocin (Sigma-Aldrich®) (40mg/kg) were administered to the HFD group, two weeks after the beginning of the diet (Yurre et al., 2020). Mice from CD group received citrate buffer 0.1M (vehicle). The body weight was registered weekly. After seven weeks of diet, mice were submited to behavioral tests and tissue collection for biochemical and cellular analisys (Figure 1A). All animal procedures fulfilled the guidelines on animal care from the NIH (National Research Council, 2011) and have been approved by the University of Brasilia Ethics Committee on the Use of Animals (CEUA/UnB, No 60/2019).

**Figure 1.**
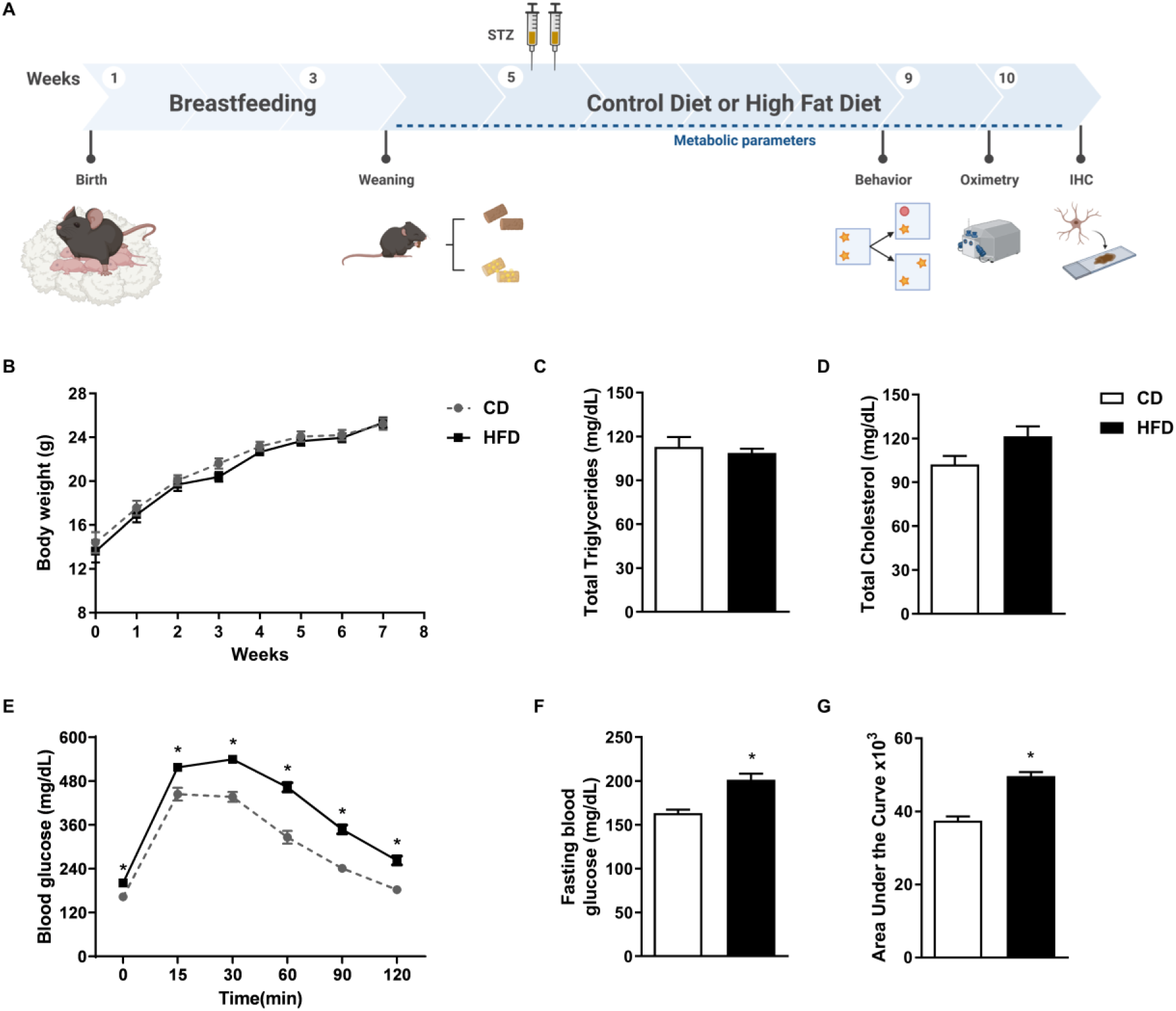
Early life HFD impaired glucose metabolism in mice. A-Experimental design: 21-day-old animals fed with either CD or HFD for 7 weeks. HFD group received 2 consecutive injections of STZ 40 mg/kg, 2 weeks after diet introduction. **B**- Weekly recorded weight from animals. **C-** Total cholesterol and **D-** triglycerides levels. **E-** Glucose tolerance test (GTT) performed after 6h of fasting (time = 0) and 15, 30, 60, 90, and 120 min after administration of glucose 2g/kg i.p. **F-** Fasting blood glucose after 6h of fasting. **G-** Area under the curve obtained from graph in Figure 1E. Cholesterol and triglycerides analysis, n= 5-7/group. Other experiments, n=22-23/group. All data were expressed as mean ± SEM. Statistical analysis was performed using student *t-test*. *p<0.05 *vs* CD.

### Behavioral Tasks

#### Novel Object Recognition (NOR) Test

This test evaluated hippocampal-dependent recognition memory (Ainge et al., 2006). NOR was performed 24 hours after preliminary 5- minute habituation in the open field arena (inside dimensions 30 x 17 x 30 cm (length x height x width). During the training session, mice were exposed for 5 min to two identical objects (Old Object, O) placed parallel to each other and 5 cm far from the walls. The animals returned to their home cages immediately after training. One hour and a half or 24 h after the training session, animals were reintroduced into the same arena for 5 minutes, with one OO and another new object (N). The arena was cleaned with ethanol 70% between each animal trial. The objects were of similar exploratory level, physical complexity, and size. The exploration time was computed when the animals touched or smelled the object at least 1 cm away. A discrimination index (DI) was calculated to evaluate recognition memory using the formula: DI = T_N_*100/(T_N_+T_O_), where T_N_ and T_O_ are the exploration time of new and old objects, respectively.

#### Object Location (OL) Test

*OL* was performed to evaluate spatial memory in rodents (Murai et al., 2007). A distinct group of animals from NOR were used in OL test. During the training session, animals were placed for 5 minutes in the arena with two identical objects located parallel to each other and 5 cm far from the walls. After the training phase, the mice were removed from the apparatus for 1.5 hours. After the inter-trial interval, one object remained in the same location (Non-displaced object, ND) and the other one was displaced to a new location (Displaced object, D). The animals were reintroduced in the arena and allowed to explore for 5 minutes. After each trial, the experimental apparatus was cleaned with ethanol 70%. The exploration time was computed as in the NOR test. A location index (LI) was calculated to evaluate location memory using the formula: LI = T_D_*100/(T_D_+T_ND_),where T_D_ and T_ND_ are the exploration time of displaced and non-displaced objects, respectively.

#### Forced swimming (FS) test

This test evaluates animal responsiveness to an inescapable stress situation (Porsolt et al., 1977). The animals were individually placed into a 5 L beaker containing water 30 cm deep, at 22° C. The animals were allowed to swim in the beaker for 6 min. Immobility was defined as floating in the water without struggling, only performing necessary movements to keep the head out of the water. The first 2 min of recording were excluded and the immobility time was analyzed in the 4 last minutes.

### Glycemic and lipid profile analysis

#### Glucose Tolerance Test

A glucose tolerance test (GTT) was performed after six hours of fasting, without water restriction. Caudal blood samples were collected to determine fasting blood glucose (time 0 min), and then the animals received an intraperitoneal glucose 2g/kg injection. Caudal blood samples were collected 15, 30, 60, 90, and 120 min after glucose load and analyzed with reactive strips in a glucosimeter (Accuchek Performa, Roche®).

#### Cholesterol (CT) and triglycerides (TG) levels

After euthanasia, blood was collected and total cholesterol and triglycerides levels were measured in plasma by colorimetric assays following manufacturers’ instructions. (LABTEST® comercial kits).

### High-resolution respirometry

Oxygen consumption was evaluated by high-resolution respirometry (HRR) in OROBOROS Oxygraph-2k at 37°C (Oroboros Instruments, Innsbruck, Austria). A subgroup of mice received ketamine and xylazine i.p. (80:8/mg/kg, respectively) until total analgesia and then were euthanized by cervical dislocation. Hippocampi, BAT, heart, and liver were collected immediately after the euthanasia.

*Hippocampal respirometry:* Hippocampi were collected and homogenized in 300 µl of isolation buffer [IB: mannitol 225 mM, sucrose 75 mM, Hepes 10 mM, EGTA 1mM, bovine serum albumin (BSA) fatty acid-free 0.1%] using a 5 mL glass-teflon homogeneizer. The samples (∼0.200 mg/mL) were added to a 2-ml chamber containing reaction buffer (RB: sucrose 125 mM, KCl 65 mM, K_2_HPO_4_ 2 mM, MgCl_2_ 1 mM, Hepes 10 mM, EGTA 0.2 mM).

Substrates (all purchased from Sigma-Aldrich) were added sequentially to assess O_2_ flux, as follows: pyruvate + malate (PM: 5 and 2.5 mM, respectively) or rotenone + succinate (ROT+S: 0.5 µM and 10 mM, respectively). State 3 (phosphorylating) was assessed after the addition of adenosine diphosphate (ADP: 500 µM), and state 4 (leak) after the addition of oligomycin (OMY: 0.1 µg/mL). Maximum respiration was evaluated after the uncoupler carbonyl cyanide 3-chlorophenylhydrazone (CCCP) titration (final concentration 1-3 µM), and non-mitochondrial residual respiration was evaluated with ROT (0.5 µM) or Antimycin A (AA, 1 µM).

*BAT respirometry:* BAT was collected, weighted and cut into 1 mm^3^ pieces. About 10x volume (µL) of RB corresponding to BAT weight was added to homogenize the tissue into a 5 mL glass-teflon homogenizer. The samples (∼0.135 mg/mL) were added to a 2 ml chamber containing RB, and then were sequentially added PM (5 and 2.5 mM, respectively), guanosine diphosphate (GDP: 1mM) to evaluate UCP-1 function, CCCP (3-7 µM final concentration), and ROT (0.5 µM).

*Liver and heart respirometry*: Tissues were collected and cut into 1mm^3^ pieces in a beaker containing IB. Successive washes were done to remove the excess of blood. Then, the tissues were homogenized into a 30 mL glass-teflon homogenizer, and it was added about 10x volume (µL) of IB according to the total amount of the corresponding tissue. The homogenates were centrifuged at 600 g. Next, the supernatant was collected and centrifuged at 7000 g. Finally, the supernatant was discarded, and the pellet was resuspended in 10 mL of IB and centrifuged at 7000 g. All centrifugations were performed at 4 °C per 10 minutes. The remaining pellet was resuspended in 500 µL of IB. The final samples were added to a 2 ml chamber containing RB, and substrates were sequentially added (PM: 5 and 2.5 mM, respectively, or ROT+S: 0.5 µM and 10 mM, respectively), ADP 500 µM, OMY 0.1 µg/mL, CCCP (final concentration 1-5 µM), ROT or AA (0.5 or 1 µM, respectively).

### Immunohistochemistry: GFAP and NeuN staining

Another subgroup of animals received ketamine and xylazine i.p. (80:8/mg/kg, respectively) until total analgesia and was intracardially perfused through the left cardiac ventricle with 0.1 M phosphate-buffered saline (PBS), pH 7.4, with the assistance of a peristaltic pump (4 mL/min). After completing the perfusion, the animals were decapitated, and their brains were removed, post-fixed in buffered PFA 4% overnight at room temperature, and cryoprotected by immersion in a 30% sucrose solution in PBS at 4°C, until complete saturation. Brains were frozen and stored at -80°C, and sliced in a cryostat into 30 µm-thick sections at -20°C.

Astrocytic proliferation was evaluated by GFAP-immunolabeling (Sofroniew and Vinters, 2010), and neuronal density was evaluated by NeuN-immunolabeling (Gusel’nikova and Korzhevskiy, 2015). Thirty µm free-floating slices were incubated with citrate buffer for 60 min at 70 °C, for antigenic recovery. Next, slices were washed with tris buffer saline (TBS) and blocked for 2 h with blocking solution (4% BSA in TBS + 0.5% triton, TBST). After that, slices were incubated overnight with primary antibody mouse anti-GFAP (1:200; Sigma-Andrich®) or rabbit anti-NeuN (1:750; Millipore®). Then, slices were washed with TBST and incubated with secondary antibody anti-mouse or anti-rabbit. Streptavidin Peroxidase and DAB immunohistochemistry kits (Polyvalent HRP/DAB detection kit for Mouse and Rabbit Specific HRP/DAB, Abcam®) were used following manufacture instructions. Slices were washed, mounted in gelatinized slides, incubated for 20s with hematoxylin (Harris hematoxylin, Merck®) and dehydrated with alcohol followed by xylene. Slides were mounted with varnish and coverslipped. Slides were scanned with Aperio Digital Pathology Slide Scanner CS2 (Leica Biosystems, Wetzlar, Germany) in 20X magnification, and pictures from hippocampal regions CA1, CA2, CA3 and DG were taken to get a representative sample of the whole hippocampus. Then, it was measured the total area stained from labeled cells using the software ImageJ® (Schindelin et al., 2012).

### Statistical analysis

All data were expressed as mean ± SEM and were analyzed with GraphPad Prism 6.0®. Kolmogorov-Smirnov normality test was performed on all samples. One-sample *t*-test was performed for NOR and OL tests to determine if the discrimination or location indexes differ from a hypothetical value of 50%. Mitochondrial respirometry parameters were analyzed by paired Student *t-test* for Gaussian distribution and Wilcoxon test for non-gaussian distributions. All other experiments were analyzed by unpaired Student *t-test* for Gaussian distribution and Mann-Whitney test for non-gaussian distribution. Results were considered statistically significant for p<0.05.

## 3. RESULTS

### Glucose metabolism is impaired in early life HFD treated mice

The introduction of an HFD early in life (P21) with 2 consecutive injections of low doses of STZ, on P35 and P36, induced changes in glucose metabolism of young adult mice. It is important to mention that STZ dosage used here does not induce massive destruction of pancreatic islet β-cell and reduction of insulin secretion as occurs with the use of higher doses (e.g., 200 mg/kg) (Furman, 2015; Wu and Yan, 2015). Seven weeks of HFD protocol did not change the body weight (Figure 1B) and lipid profile, evaluated by plasma cholesterol and triglicerides levels (Figure 1C-D) in HFD mice. Although, glucose uptake was severely impaired (Figure 1E) and fasting blood glucose levels were significaly increased in HFD mice (Figure 1F). We observed an increase in glucose levels in all of the six time points, with glucose levels remaining elevated even 120 min after glucose load (Figure 1E). Furthermore, the area under the curve of glucose tolerance test was higher in HFD mice (Figure 1G).

### Hippocampal-dependent memory is impaired in early life HFD-fed mice

Regardless of the experimental group, mice showed no differences in baseline exploratory activity during the habituation period in the open field arena (Figure S1 A-C). CD animals demonstrated typical short (1.5h) or long-term (24h) recognition memories, as shown by increased exploratory behavior towards novel objects used in test sessions. However, HFD mice did not reach the exploratory percentage of the novel object above the hypothetical value of 50%, 1.5 h after training section (Figure 2A). With a 24h delay, HFD mice discriminated the novel object, but the exploration time from these animals was significantly lower than CD ones (Figure 2B). Regarding spatial memory, only the CD group demonstrated a location index significantly above the hypothetical value, while the HFD mice did not discriminate the object’s new position (Figure 2C). Also, in the evaluation of depressive-like behavior through the FS test, there were no differences in immobility time between HFD and CD groups (Figure 2D).

**Figure 2.**
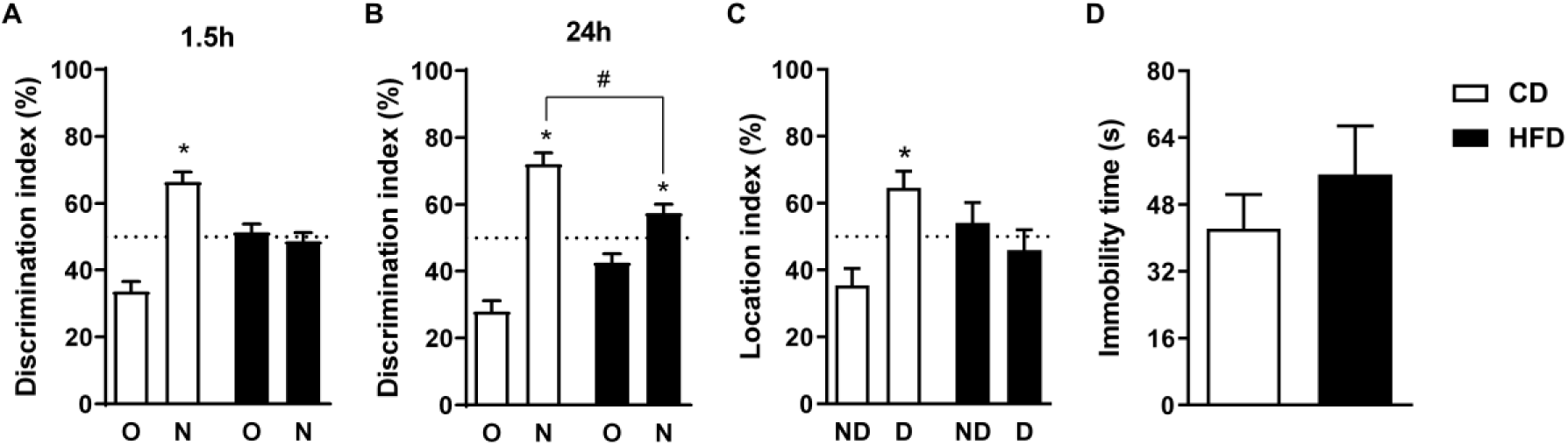
Early life HFD affected cognitive function in mice. Cognitive performance was evaluated by the discrimination index, accessed by NOR using a **A**- 1.5 h and a **B**- 24 h as intersection intervals, and by **C-** location index, accessed by OL using a 1.5-hour intersection interval. **D-** Immobility time obtained from FS test. NOR and FS, n= 15-16/group. OL, n=7/group. O = old object, N = new object, ND = non-displaced object, D = displaced object. All data were expressed as mean ± SEM. Statistical analysis was performed using student *t- test*. *p<0.05 *vs* 50% chance levels, ^#^p<0.05 *vs* N of CD group.

### Early life HFD caused hippocampal mitochondrial dysfunction and astrogliosis in mice

HFD mice presented decreased hippocampal oxygen consumption on state 3 and state 4 U (uncoupled) in complex-1 and 2 driven respiration (Figure 3A, B, E, F). These impairments affected the ATP production (Figure 3C, G) and the respiratory reserve capacity in complex-1 driven respiration (Figure 3D), with no changes on respiratory reserve capacity using complex- 2 linked substrates (Figure 3H). Also, there was no difference in safranine fluorescence between the experimental groups (Figure S2), indicating that the H^+^ gradient generated during electron flow through the respiratory chain was preserved.

**Figure 3.**
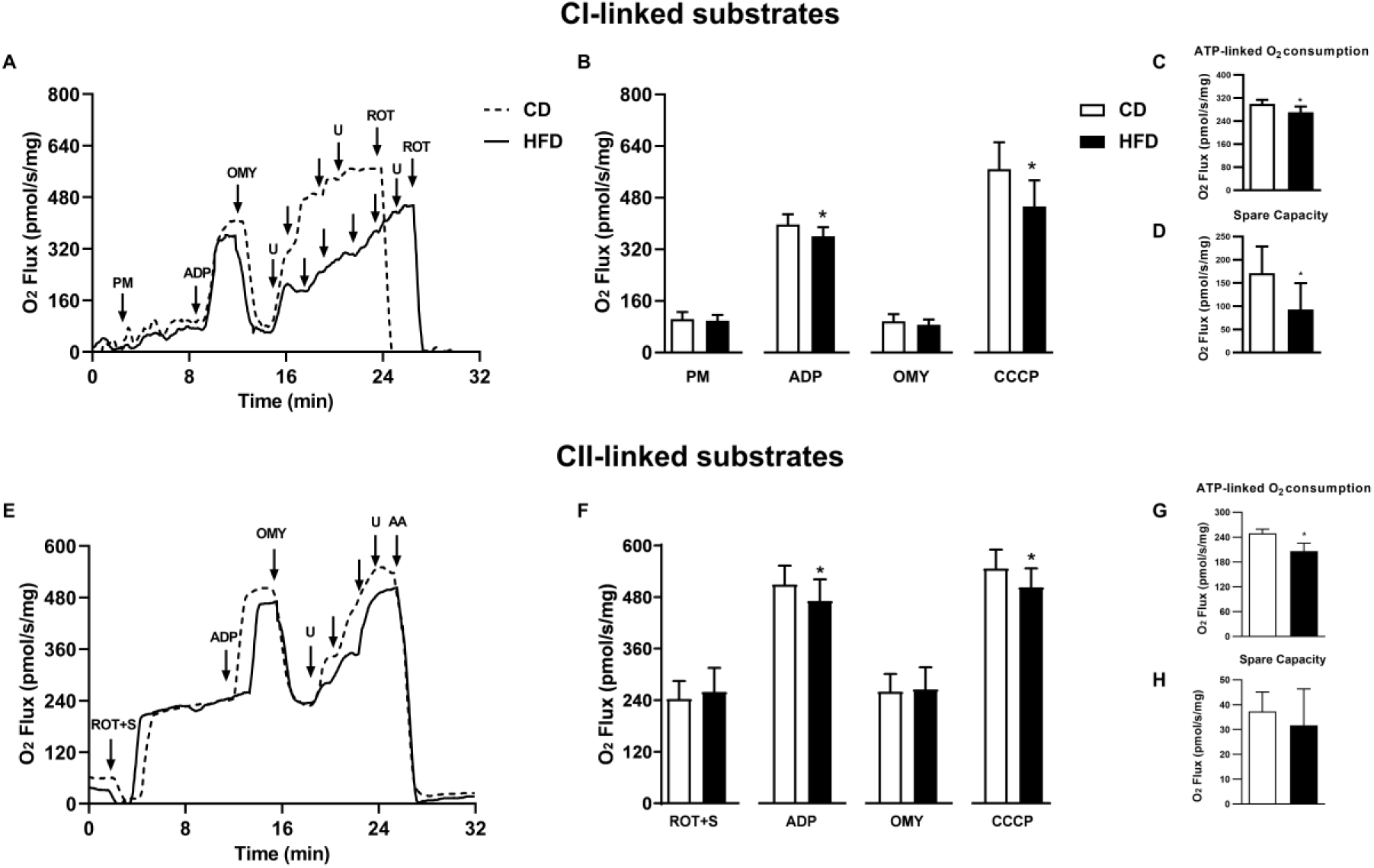
Early life HFD impacts hippocampal oxygen consumption. HRR was performed to evaluate mitochondrial respiratory states: State 2 (PM or ROT+S), state 3 (ADP), state 4 (OMY) and state 4 uncoupled (U=CCCP). **A-D**: results from complex 1-linked substrates and **E-H** results from complex 2-linked substrates. **A** and **E**- representative oxygraphs from complex 1 and complex 2 linked substrates, respectively. **B** and **F**- O_2_ flux (pmol/s) *per* mg of protein in all respiratory states. **C** and **G**- ATP-linked oxygen consumption, calculated by the difference between state 3 and state 4. **D** and **H-** Spare capacity, calculated by the difference between state 4 uncoupled and state 3. Complex 1-linked substrates: n=6/group. Complex 2- linked substrates: n=8/group. All data were expressed as mean ± SEM. Statistical analysis was performed using student *t-test*. *p<0.05 *vs* CD group.

The quantification of total labeled cells in the whole hippocampus was done by the sum of labeled cells in CA1, CA2, CA3 and DG regions (Figure 4A). Astrogliosis, here evaluated througth GFAP labeling, was significantly increased in hippocampal regions CA1, CA3 and DG, without significant changes in CA2 region (Figure S3 A-D). The sum of labeled GFAP cells in the whole hippocampus was significantly increased in HFD mice compared to CD ones (Figure 4C). Moreover, we found a decrease in NeuN staining in hippocampal CA1 and CA2 regions from HFD mice, with no changes in CA3 and DG (Figure S3 E-H). A trend towards a decrease (p=0.0762) in NeuN staining in the whole hippocampus of HFD mice was observed (Figure 4E).

**Figure 4.**
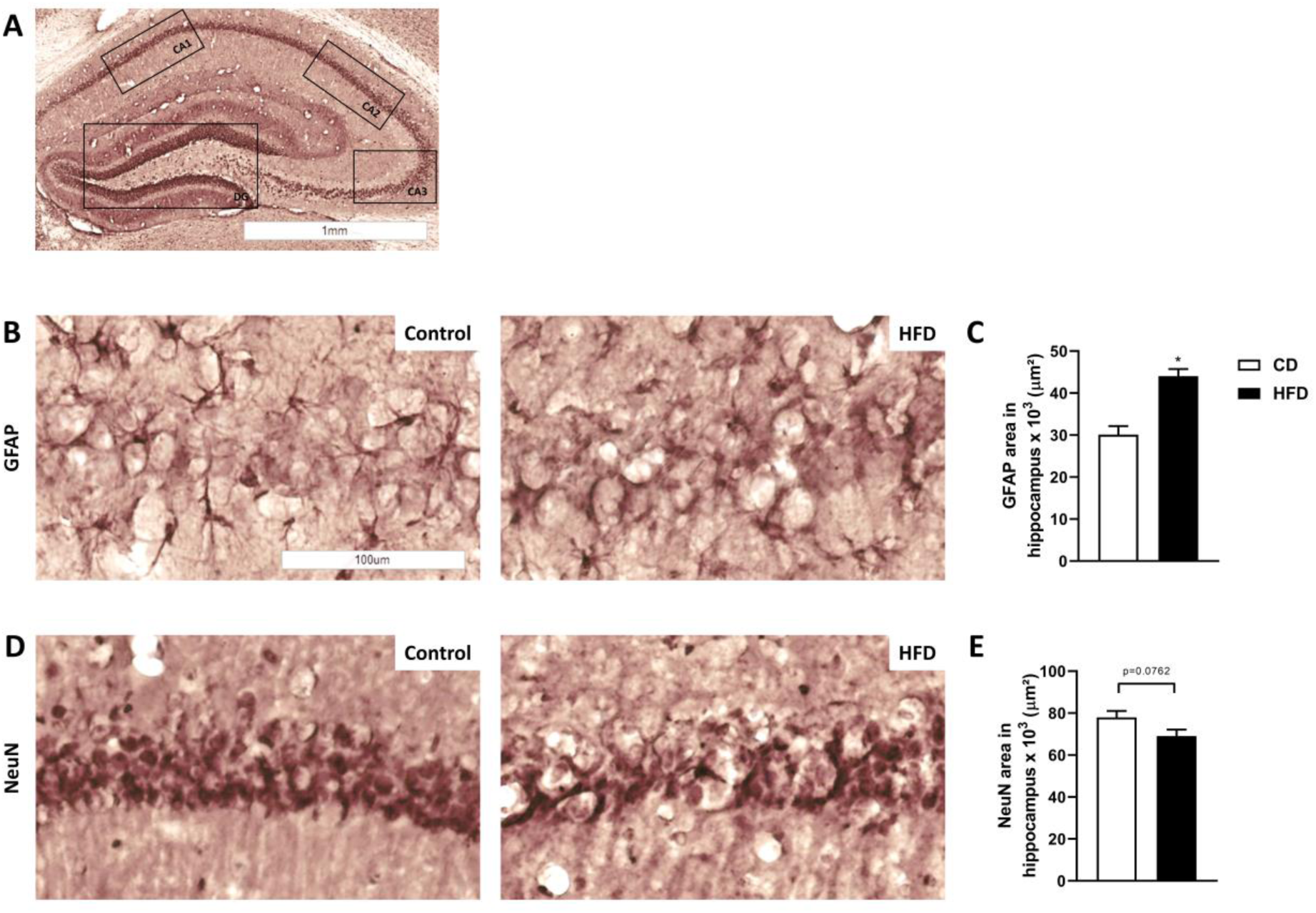
Effect of early life HFD in hippocampal cellularity. A-. Representative hippocampal slices indicating the regions CA1, CA2, CA3 and DG, where the quantification of labeled cells was done. **B** Representative images of GFAP staining in CA1 region, and **C-** quantification of total GFAP stained area in the whole hippocampus. **D** Representative images of NeuN staining in CA1 region, and **E-** quantification of total NeuN stained area in the whole hippocampus. n=5/group. All data were expressed as mean ± SEM. Statistical analysis was performed using student *t-test*. *p<0.05 *vs* CD group.

### Early life HFD promoted mitochondrial dysfunciont in the BAT and liver, but not in the cardiac tissue

To investigate the effect of early life HFD in peripheral tissues, we performed HRR in BAT homogenates and in isolated mitochondria from liver and heart. BAT homogenates from HFD mice, energized with PM, showed reduced O_2_ consumption when compared to CD mice (Figure 5A, B). As expected, BAT oxygen consumption from CD mice was strongly inhibited after GDP addition, as a result of UCP-1 inhibition. The same pattern was not observed in BAT homogenates from HFD mice (Figure 5A,B). The UCP-1 linked respiration was significantly reduced in HFD mice (Figure 5C). However, when challenged with the uncoupler CCCP, BAT mitochondria from this group presented an enhanced respiratory capacity when compared to CD mice (Figure 5D). Therefore, BAT mitochondria from HFD mice may work far from their maximum capacity (around 50%), while these mitochondria from CD mice work at their full capacity.

**Figure 5.**
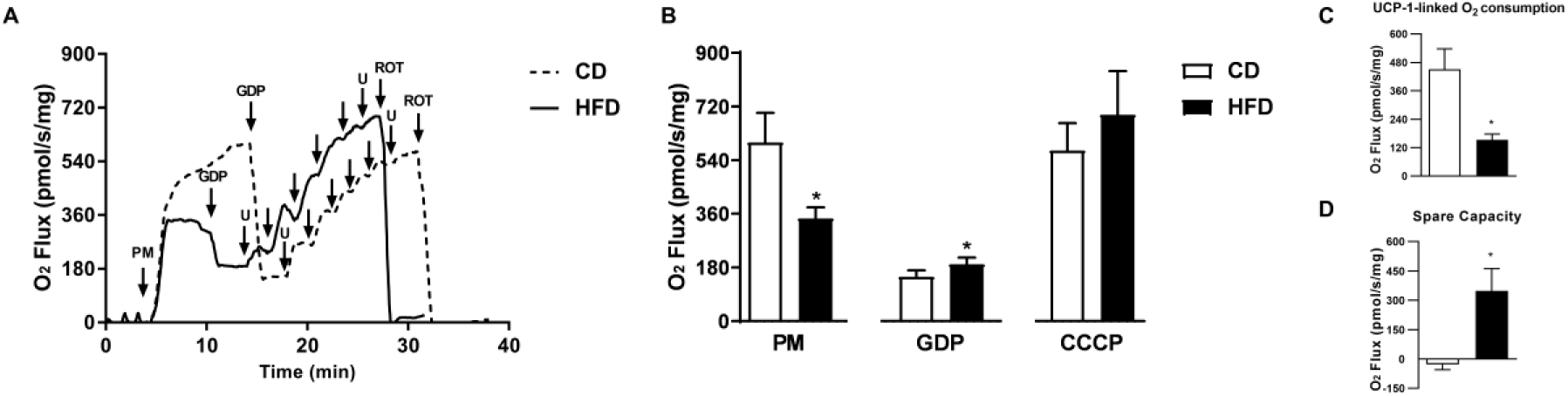
Early life HFD impacts BAT oxygen consumption. High-resolution respirometry performed to evaluate mitochondrial respiratory states: State 2 (PM), UCP-independent respiration (GDP) and state 4 uncoupled (U=CCCP). **A-** Representative oxygraphs from respiratory states. **B-** O_2_ flux (pmol/s) *per* mg of protein in all respiratory states. **C-** UCP linked- respiration calculated by the difference between state 2 and UCP-independent respiration. **D-** Spare capacity calculated by the difference between state 4 uncoupled and state 2. n=6/group. All data were expressed as mean ± SEM. Statistical analysis was performed using student *t- test*. *p<0.05 *vs* CD group.

In liver, complex-1 driven respiration was decreased on states 2 and 4 in HFD mice (Figure 6A). However, these effects were not maintained when liver mitochondria were energized with complex-2 linked substrates (Figure 6B). Also, safranine fluorescence was not different between the experimental groups (Figure S4A). No changes were verified in heart mitochondria oxygen consumption of HFD mice using neither complex-1 nor complex 2 linked substrates (Figure 6C,D), as well as, no significant alteration was observed in safranine fluorescence (Figure S4B).

**Figure 6.**
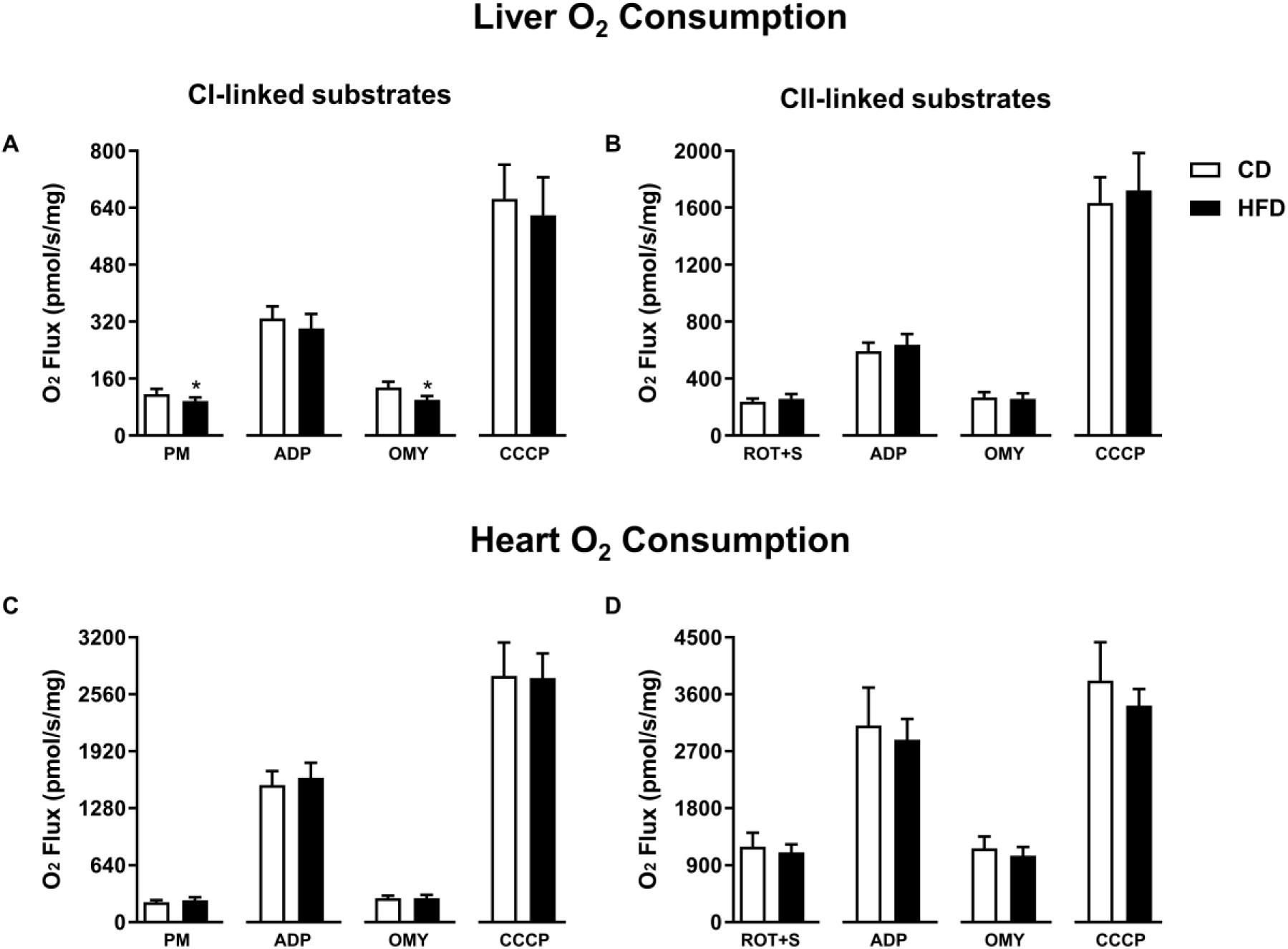
Effect of early life HFD on liver and heart oxygen consumption. High-resolution respirometry performed to evaluate mitochondrial respiratory states: State 2 (PM or ROT+S), state 3 (ADP), state 4 (OMY) and state 4 uncoupled (U=CCCP). Liver O_2_ flux (pmol/s) *per* mg of all respiratory states from **A-** Complex-1 and **B-** Complex-2 linked substrates. Heart O_2_ flux (pmol/s) per mg of protein in all respiratory states from **C-** Complex-1 and **D-** Complex-2 linked substrates. Liver complex 1-linked substrates: n=7/group. Liver complex 2-linked substrates: n=8 /group. Heart complex 1-linked substrates: n=9/group. Heart complex 2-linked substrates: n=6/group. All data were expressed as mean ± SEM. Statistical analysis was performed using student *t-test*. *p<0.05 *vs* CD group.

## 4. DISCUSSION

Herein, we showed that feeding juvenile mice with HFD from weaning to adulthood, induce memory impairment and hippocampal dysfunction, as well as bioenergetic dysfunction in metabolic active tissues, leading to glucose intolerance, peripheral (BAT and liver) and central (hippocampus) mitochondrial dysfunction, despite not inducing overweight/obesity and changes in lipid profile. We used a combination of HFD with low STZ dose in order to increase the metabolic disturbance, consistent with high glucose levels and insulin resistance described elsewhere (Yurre et al., 2020). Similar dietary intervation starting from weaning also induced metabolic changes, such as dyslipidemia, hyperglycemia, and changes in leptin and insulin levels, in absence of obesity in juvenile mice (Boitard et al., 2012; Vinuesa et al., 2016).

Metabolic disturbances were accompanied by impairment in short and long-term recognition and spatial memory indexes in HFD juvenile mice, without inducing depressive-like behavior. Our data highlight phenotypic hippocampal changes that are usual consequences of functional and structural dysfunction. The process of memory formation relies on synaptic plasticity, in which many receptors and intracellular signaling pathways are involved in structural and functional neuronal changes, that induce and maintain the long-term potentiation (LTP), which is the basis of memory formation (Bliss and Cooke, 2011). Using 1.5 h as intersection interval in the NOR and OL tests, we evaluated a short-term memory, which is associated with an early LTP, where there is enhanced conductance and abundance of glutamatergic receptors activation in postsynaptic neurons. Regarding long-term memories, it is believed that gene expression and protein synthesis take part in the generation of new AMPA receptors, as well as novel dendritic spines and synaptic connections, which are associated with the NOR test 24h intersection interval used in our study (Baltaci et al., 2019). Short and long-term memory impairment were here observed using NOR and OL tests as recognition and spatial cognitive assessments, respectively. Our data are in agreement with recent studies that deescribed cognitive dysfunction in juvenile mice fed a HFD for one week (Khazen et al., 2019), 6 to 8 weeks (Vinuesa et al., 2019; Yang et al., 2019) and 12 weeks (Boitard et al., 2012; Vinuesa et al., 2016).

A recent study from our group reported memory dysfunction in adult *Swiss* mice as early as 3 days after HFD introduction (de Paula et al., 2021). Interestingly, some studies emphasize that behavior and metabolic changes are more intense when HFD is introduced earlier in life. In this context, it has been previously shown a spatial memory dysfunction and diminished LTP in juvenile, but not adult mice exposed to a one week of HFD (Khazen et al., 2019) and impairment in spatial memory with increased hippocampal cytokines in juvenile, but not adult rats, after 2 month of HFD (Boitard et al., 2014). The same group showed that only juvenile mice displayed spatial learning and relational memory impairment after 3 weeks of HFD (Boitard et al., 2012). These data reinforce our findings, which show that exposure to HFD early in life cause deep impacts on rodents behavior and metabolism, permanently compromising brain functions.

Among the metabolic deregulation caused by HFD, mitochondrial dysfunction seems to be a common feature affecting not only peripheric metabolic organs but also the brain. Specially in the hippocampus, HFD impairs mitochondrial oxidative status (Gao et al., 2019; Pintana et al., 2013), bioenergetics (de Paula et al., 2021; Wang et al., 2015), dynamics (Ruegsegger et al., 2019), biogenesis (Petrov et al., 2015) and calcium homeostasis (Park et al., 2018; Petrov et al., 2015) in adult mice. Our data bring new evidence about the correlation between mitochondrial dysfunction and behavioral changes induced by HFD early in life. We found a bioenergetic disturbance in hippocampus of juvenile HFD mice characterized by a decrease in oxygen consumption on state 3 and maximum respiration, that affected the ATP production and the respiratory reserve capacity. Also, Yang and colleagues observed changes in mitochondrial biogenesis and dynamics in hippocampus, together with depressive-like behavior and spatial memory impairment in adult mice chronically treated with a HFD (Yang et al., 2021). Beside, we previously reported mitochondrial dysfunction in hippocampus of adult swiss mice treated with HFD (de Paula et al., 2021). Therefore, our data raise the flag to a possible metabolic shift in the hippocampus of juvenile HFD mice, switching to a more glycolytic and less oxidative profile. Supporting this hypothesis, a study from Wang and colleagues (2020) reported an increased activity of fatty acid synthase and glycolytic enzymes in the hippocampus, associated with oxidative stress and LTP impairment in HFD adult mice. (Wang et al., 2020).

Indeed, mitochondrial impairment impacts cellular function in the hippocampus. This puts mitochondria (dys)function as one of the key elements involved in structural and functional hippocampal changes caused by HFD, making the environment more inflammatory and auspicious to dysfunctional synapsis (Dutheil et al., 2016; Hao et al., 2016). In accordance with this, Park and colleagues demonstrated that long-term HFD associated with STZ caused neuroinflammation and impaired hippocampal mitochondria homeostasis by increasing mitophagy and reducing the number of synapses, dendritic spines and also the number of presynaptic mitochondria in adult mice (Park et al., 2021). Bioenergetic dysfunction may be the center of these effects since lower neuronal ATP concentrations trigger hyper-reactivity of astrocytes in order to restore the neuronal ATP levels (Garcia-Serrano and Duarte, 2020).

It is also reported the relation between hypercaloric diet-induced metabolic syndrome/insulin resistance, and hippocampal inflammation, astrogliosis, oxidative stress, and memory impairment (Trevino et al., 2015; Trevino et al., 2017). Here we found an increase in GFAP staining in the hippocampus of juvenile HFD mice. Since astrocytes have a key role in cerebral homeostasis, changes in their morphology and function may directly impact their perisynaptic and endfeet process, essential to regulate neurotransmitter, cytokine and growth factor levels, as well as to preserve cerebral blood flow and blood-brain barrier homeostasis (Belanger et al., 2011). Accordingly, other studies showed hippocampal glial activation in juvenile HFD fed mice, with focus on increasing microglia activation, inflammatory cytokines production (Boitard et al., 2014; Vinuesa et al., 2019; Vinuesa et al., 2016; Yang et al., 2019) and decreased neurogenesis (Boitard et al., 2012; Vinuesa et al., 2016). Although we did not observe a significant difference in neuronal density across the hippocampus, there was a decrease in NeuN staining specifically in CA1 and CA2 regions of juvenile HFD mice. In line with this, a recent study correlated the negative effects of HFD feeding with neuronal synaptic damage, leading to impaired synaptic plasticity (Crispino et al., 2020).

Furthermore, we should not rule out hyperglycemia effects on glutamatergic hippocampal homeostasis, and its close relation to memory loss. Ripoli and colleagues reported a reduced hippocampal-mediated firing mediated by AMPA and NMDA, and altered subunit content (GluR1, GluN and GluA) in STZ-induced hyperglycemia in mice (Ripoli et al., 2020). These alterations are strongly correlated with LTP reduction and cognitive impairment (Ripoli et al., 2020; Viswaprakash et al., 2015). Also, changes in hippocampal brain-derived neurotrophic factor (BDNF) may contribute to memory loss, since reductions in its levels are associated with recognition (Ripoli et al., 2020) and spatial memory dysfunction (Stranahan et al., 2008). Since the hippocampus is essential in the process of memory formation, and especially in spatial memory establishment (Barker and Warburton, 2011), it is very plausible that the memory impairment observed in our study might be due to hippocampal neuronal dysfunction, facilitated or exacerbated by astrogliosis and mitochondrial bioenergetics dysfunction.

Other molecules produced in peripheral tissues, and not only within the SNC, are able to modulate cognitive processes. Despite the primary function of brown adipocytes is to dissipate energy as heat via UCP-1, BAT also has a critical role in modulating metabolism as well via *“batokines”* and microRNAs (Lee et al., 2019; Villarroya et al., 2017). For example, interleukin-6 (IL-6), a mediator of inflammation and immune response, is essential for spatial learning (Bialuk et al., 2018) and recognition memory (Baier et al., 2009; Bialuk et al., 2019). IL-6 derived from BAT is necessary to control glucose homeostasis, weight gain, adiposity, and FGF21 levels (Stanford et al., 2013). Interestingly, FGF21 secreted from the liver and BAT improves hippocampal neurogenesis, inflammation, and metabolic parameters, all related to recognition memory and anxiety-like behavior in obese-insulin resistant mice (Wang et al., 2018).

BAT mitochondrial dysfunction were observed in our juvenile HFD model and suggests a *whitening* process and a metabolic shift, since oxidative UCP-1-uncoupled activity, was diminished. In accordance, previous studies reported that BAT hyperthrophic lipid droplets and macrophage infiltration (Dinh et al., 2015; Gao et al., 2020), decreased substrate oxidation and UCP-1-linked oxygen consumption (Pajuelo et al., 2012), impaired mitochondrial function, dynamics and autophagy, as well as decreased noradrenergic stimuli into BAT and brainstream (Silvester et al., 2018) in HFD-treated adult rodents.

Hepatic alterations are very recurrent in HFD rodent models (Lv et al., 2019; Mantena et al., 2009) and in overweight/obese humans (Koliaki et al., 2015). Liver mitochondrial dysfunction is a common feature of HFD, and it is believed that a reduced hepatic oxidative capacity contribute to steatosis development (Meex and Watt, 2017), however there is little evidence on the early-life effects of HFD on hepatic mitochondrial function. The juvenile mice also displayed impaired hepatic bioenergetic function after receiving HFD. Hepatic inflammation, elevated lipid content and diminished insulin signaling were previously observed in juvenile mice exposed to a 16-week HFD (Vinuesa et al., 2016), as well as steatosis, inflammation, increased lipid metabolism and changes in microbiota profile in weaned mice exposed to a 11-week HFD (Carbajo-Pescador et al., 2019). Some of the mitochondrial changes observed in the liver of hyperglycemic patients include lower levels of inorganic phosphate and ATP, and reduced ATP turnover (Schmid et al., 2011; Szendroedi et al., 2009), while patients with liver disease presented reduced mitochondrial mass and biogenesis factors, as well as oxidative damage (Koliaki et al., 2015).

Adult mice with either HFD+STZ or just HFD-induced obesity present diminished liver activity of complexes I, II, IV, V, associated with increased oxidative stress and LDH activity in liver, skeletal muscle, and WAT (Jha and Mitra Mazumder, 2019). Additionally, we observed liver dysfunction, without state 3 respiration changes, is in accordance with previously published data, which reported reduced ATP-linked respiration in adult mice only after at least 16 weeks of HFD (Mantena et al., 2009). Thus, HFD may induce a multi-organ metabolic shift to the glycolytic pathway, reducing mitochondrial oxidative phosphorylation efficiency, as well as increasing lactate, which triggers inflammation and tissue damage (Jha and Mitra Mazumder, 2019).

Interestingly, our data did not indicate any cardiac bioenergetic change in juvenile HFD mice. Adult mice submitted to same hyperglycemic-HFD model, presented changes in cardiac mitochondrial function after normalizing the O_2_ consumption by the citrate synthase activity, a notable marker of mitochondrial content, suggesting that the cardiac bioenergetic alterations were associated with reduced mitochondrial content, not mitochondrial dysfunction (Yurre et al., 2020).

Together our data show that exposing weaned mice to HFD induced a multi-organ metabolic shift, impacting mitochondrial oxidative phosphorylation efficiency, which in turn can trigger inflammation and tissue damage. BAT was the most affected peripheral tissue, presenting a bioenergetic evidence of “whitening”, that may worsen systemic glucose metabolism and inflammation. On the CNS, hippocampal function and cell density were altered, downregulating mitochondrial electron transport chain, which may have triggered reactivity of astrocytes, neuronal loss, and cognitive impairment (Figure 7). Central and peripheral metabolic alterations may be connected and may occur even before obesity itself. Our data open a new perspective to explain the metabolic and behavioural outcomes triggered by introducing a HFD early in life, focusing in the establishment of a general bioenergetic crisis.

## Funding

This work was supported by the Fundação de Apoio à Pesquisa do Distrito Federal (FAPDF grants 00193-00000229/2021-21, 00193-00001324/2019-27); Conselho Nacional de Desenvolvimento Científico e Tecnológico (CNPq grant 424809-2018-4); Coordenação de Aperfeiçoamento de Pessoal de Nível Superior (CAPES/STINT grant 88881.465507/2019-01); and Instituto Nacional de Ciência e Tecnologia e Neuro-ImunoModulação (INCT-NIM grant 485489/2014-1).

## Acknowledgments

We gratefully acknowledge Prof. Emiliano Horácio Medei from Carlos Chagas Filho Biophysics Institute, Federal University of Rio de Janeiro, for the discussions and suggestions on the experimental protocol, and to Priscila Batista da Rosa for the graphical design.

## SUPPLEMENTARY MATERIAL

### Methods

*Open Field (OF):* The OF was used to evaluate the locomotor and exploratory activities induced by a novel environment (Prut and Belzung, 2003). The apparatus was an arena with dimensions length x height x width of 30 cm x 30 cm x 17 cm, respectively, and a white background, containing spatial cues on the sidewalls. Animals were individually placed in the center of the apparatus to explore the arena for 5 minutes freely. The arena was cleaned with ethanol 70% between each animal trial. Total distance traveled and time spent in the internal and external quadrants (inner quadrant drew 7.50 cm away from the walls) were evaluated on ANY-maze® software (Stoelting Co., IL US).

*Mitochondrial membrane potential*: In complex 2-linked substrate experiments (ROT+S), membrane potential was estimated simultaneously with high resolution respirometry. The estimated membrane potential was determined using a fluorescence sensor (O2k-Fluorescence LED2 module, Oroboros Instrument, Austria, excitation 495 nm, emission 587 nm). The potentiometric probe Safranine (2µM) has affinity for membranes, making it useful for mitochondrial membrane potential assessment. Fluorescence measurements from samples were recorded every 2 seconds, concomitantly with O_2_ consumption recording. The safranine fluorescence was presented as “Δ*F*/*F*”, where F is the maximum fluorescence on state 4U (CCCP), and Δ*F* is F minus the steady-state fluorescence in each respiratory state.

---

**Table S1.** Nutrient formulation of the standard (CD) and high fat (HFD) diets. Caloric amount corresponding to 1kg of food (PRAGSOLUÇÕES Biociências®).

| *Component* | *Control Diet (CD)* |  | *High-fat Diet (HFD)* |  |
| --- | --- | --- | --- | --- |
|  | Kcal | %kcal | Kcal | %kcal |
| *Starch* | 2118 | 52.95 | 1326 | 26.57 |
| *Casein* | 800 | 20 | 800 | 16.03 |
| *Sucrose* | 400 | 10 | 400 | 8.02 |
| *Soy oil* | 630 | 15.75 | 360 | 7.21 |
| *Vitamin mix AIN-93* | 40 | 1 | 40 | 0.8 |
| *L-Cistin* | 12 | 0.3 | 12 | 0.24 |
| *Lard* | 0 | 0 | 2052 | 41.12 |
| *Choline* | 0 | 0 | 0 | 0 |
| *Fiber* | 0 | 0 | 0 | 0 |
| *Total* | 4000 | 100 | 4990 | 100 |

**Figure S1.**
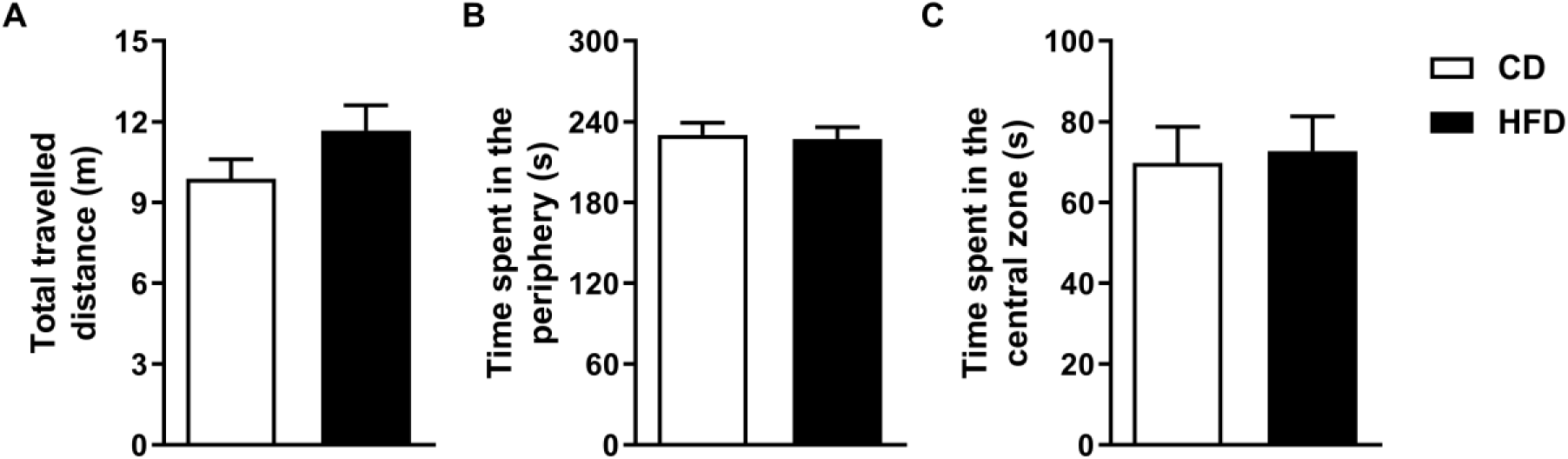
Early life HFD impact in locomotor activity. A-Total distance traveled and time spent in the B- internal and C- external zones in the open field arena, during 5 min. n=15- 16/group. All data were expressed as mean ± SEM. Statistical analysis was performed using student *t-test*.

**Figure S2.**
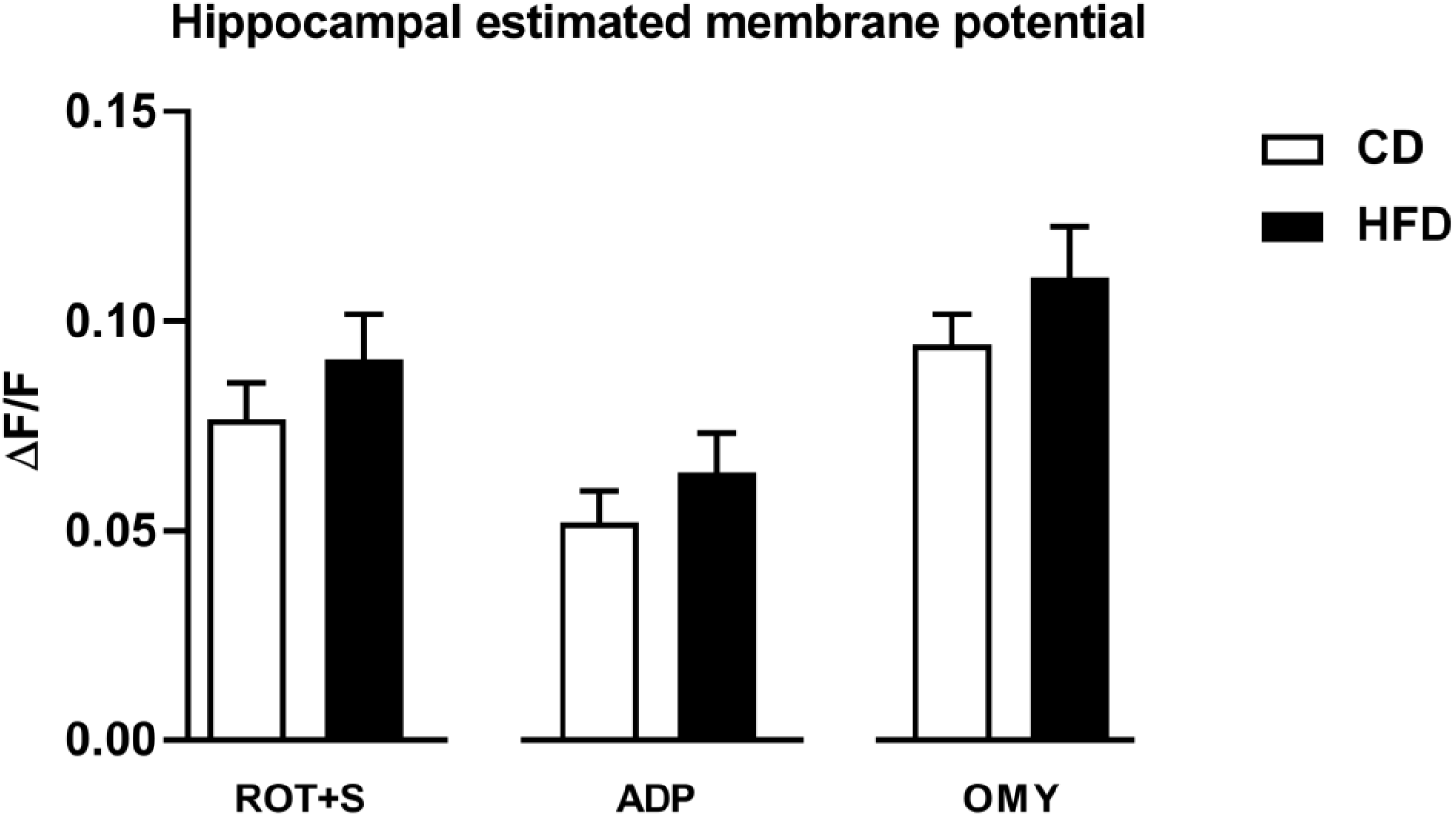
Hippocampal homogenates membrane potential. Results obtained from hippocampal homogenate membrane potential estimated by safranine fluorescence. n=8/ group. All data were expressed as mean ± SEM. Statistical analysis was performed using student *t-test*.

**Figure S3.**
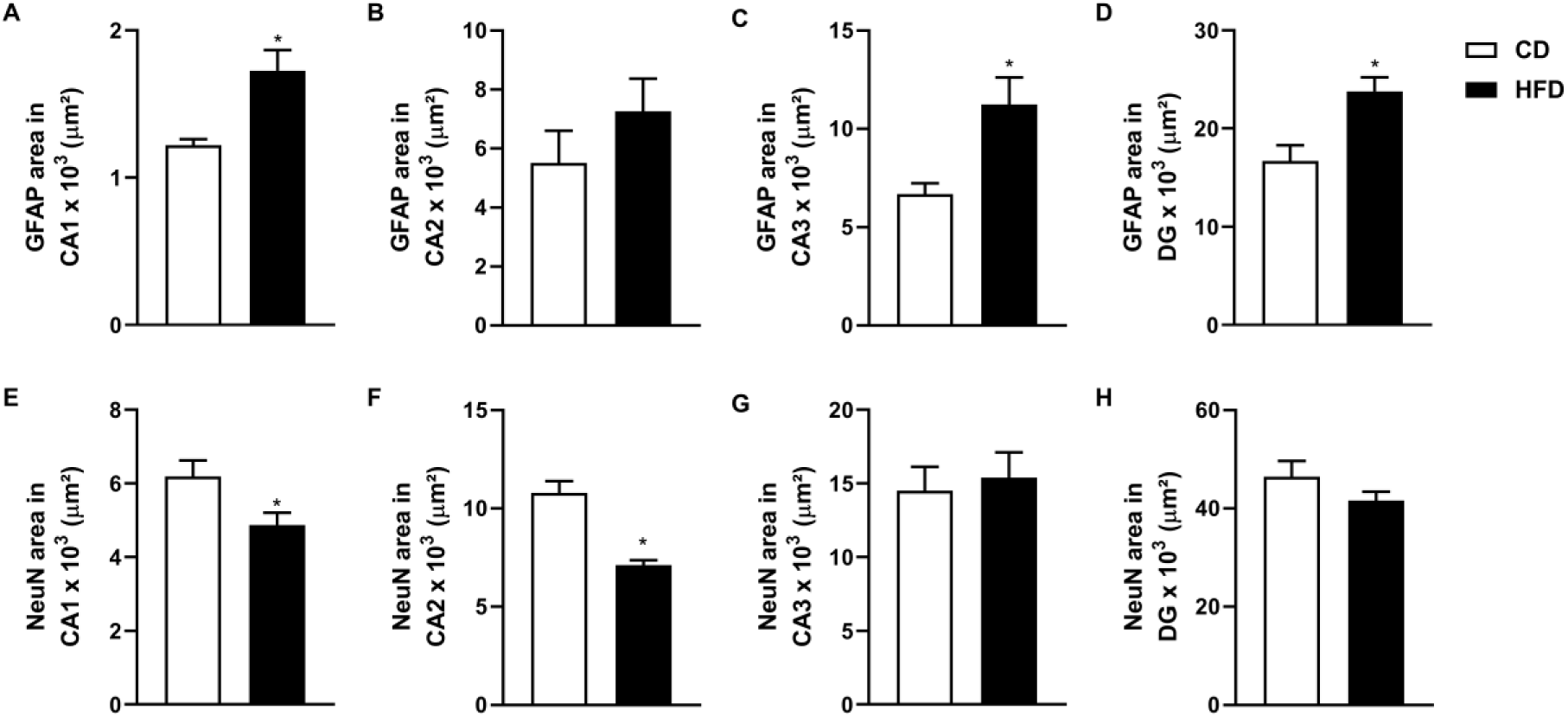
Effect of early life HFD in hippocampal regions cellularity. Quantification of total GFAP stained area in A- CA1, B- CA2, C- CA3 and D- DG regions of hippocampus. Quantification of total NeuN stained area in E- CA1, F- CA2, G- CA3 and H- DG regions of hippocampus. n=5/group. All data were expressed as mean ± SEM. Statistical analysis was performed using student *t-test*. *p<0.05 *vs* CD group.

**Figure S4.**
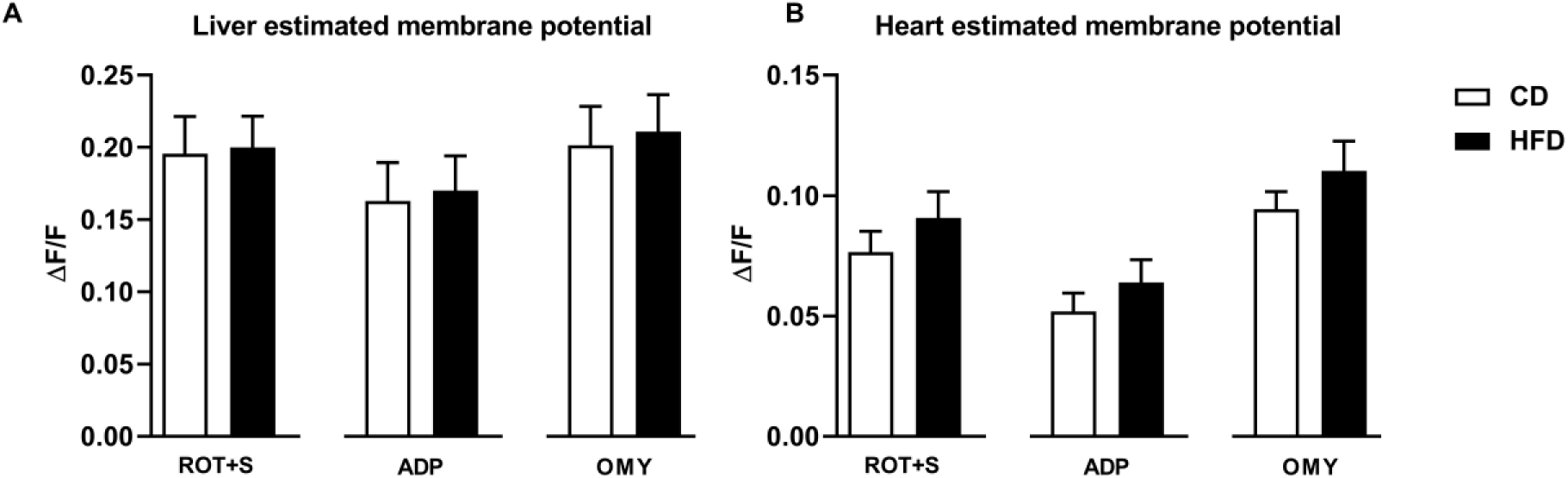
Liver and heart mitochondria membrane potential. Results obtained from A- hepatic and B- cardiac isolated mitochondria membrane potential estimated by safranine fluorescence. Liver: n=8/group. Heart: n=6/group. All data were expressed as mean ± SEM. Statistical analysis was performed using student *t-test*.

## Notes

### Competing Interest Statement

The authors have declared no competing interest.

